## Supplemental figures for "The ATM-E6AP-MASTL axis mediates DNA damage checkpoint recovery"

**Figure 1 Supplemental 1. MASTL upregulation after DNA damage.**

(A, B) SCC38 cells were treated with 10 mM HU (A) or 10 Gy IR (B), and incubated for 0-10 hours, as indicated. The expression levels of MASTL, RPA, and α-tubulin were analyzed by immunoblotting. (C, D) HeLa cells were treated with 0.5 uM DOX (C) or 10 mM HU (D) as indicated, cell lysates were collected and analyzed by immunoblotting for MASTL and α-tubulin. (E) Quantification of MASTL expression, measured by immunoblotting, and normalized to a loading control (α-tubulin or β-actin) in HeLa cells treated without or with IR, DOX, or HU. Standard derivations were calculated from three independent experiments.

**Figure 1 Supplemental 2. The expression levels of cell cycle kinases after DNA damage.**

HeLa (A) or SCC38 (B) cells were treated without or with 2 mM of hydroxyurea (HU) for 8 hours (A), or 3-6 hours (B). Cell lysates were harvested for immunoblotting.

**Figure 2 Supplemental 1. Increased protein stability of MASTL after DNA damage.** (A-C) HeLa cells expressing CFP-MASTL were treated without or with IR. Cells were analyzed by direct fluorescence for CFP-MASTL expression, and by immunofluorescence for the phosphorylation of ATM/ATR substrates. Quantifications are shown in panels B and C (>10 cells/time point). (D, E) HEK293 cells were treated without (D) or with (E) 10 mM HU for 2 hours. These cells were then treated with cycloheximide (CHX, 20 μg/ml) at time 0 to block protein synthesis, and analyzed by immunoblotting for the protein stability of MASTL and α-tubulin.

**Figure 3 Supplemental 1. E6AP and MASTL associate via their N-terminal motifs.** (A) MBP-E6AP and GST-MASTL were expressed and purified, as described in Materials and Methods. GST-MASTL on glutathione beads, or control glutathione beads, were incubated with MBP-E6AP, followed by pulldown, as described in Materials and Methods. The input, GST-MASTL pulldown, and control pull down samples were analyzed by immunoblotting for MASTL and E6AP. (B) GFP-MASTL, FL or N-terminus (aa 1-340), was expressed in HeLa cells. Immunoprecipitation (IP) was performed using a GFP antibody, and the presence of E6AP in the IP products was examined by immunoblotting. (C) IP was performed using GFP-tagged segments of E6AP expressed in HeLa cells. The IP product, cell lysate input, and a control IP (using empty beads) were analyzed by immunoblotting for MASTL and GFP. N: aa 1-280; N1: aa 1-99; N2: aa 100-207; N3: aa 108-280.

**Figure 4 Supplemental 1. E6AP mediates MASTL ubiquitination and degradation.**

(A) HeLa cells were treated without or with E6AP siRNA. The RNA levels of E6AP and MASTL were quantified by real-time PCR. The mean values were calculated from three experiments. Statistical significance was determined using an unpaired 2-tailed Student's t test. A p-value more than 0.05 was considered non-significant (ns). (B) Control or E6AP KO HeLa cells were treated without or with MG132, as indicated. The protein levels of MASTL and $\alpha$-tubulin are shown by immunoblotting. (C) The E6AP *in vitro* ubiquitination assay was performed using MASTL as substrates, as in Fig. 4H. S5a was added as a control substrate. E1/2 and E3 (E6AP) enzymes were added in the reactions, as indicated. Samples were analyzed by immunoblotting for MASTL and S5a after 90 min incubation. (D) *In vitro* ubiquitination assay was performed using E6AP as E3 ligase, and $\Delta$N MASTL (aa 335-887) as substrate. S5a was added as a control substrate. The reactions were incubated as indicated, as analyzed by immunoblotting for MASTL and S5a.

**Figure 5 Supplemental 1. Impaired DNA damage checkpoint signaling in E6AP-null cells.**

(A) Control or MASTL knockdown HeLa cells were maintained in cell culture, and cell viability was determined by counting cell numbers. The cell numbers in day 2 and 3 were normalized to those in day 1. The mean values and standard derivations, calculated from three experiments, are shown. (B) Control or E6AP knockout (KO) HeLa cells were treated with 0.1 μM etoposide (ETO) for the indicated hours. Cells were analyzed by immunoblotting for phospho-ATM/ATR substrates and α-tubulin.

**Figure 7 Supplemental 1. DNA damage-induced E6AP Ser-218 phosphorylation is mediated by ATM.**

(A) WT or E6AP knockout HEK293 cells were treated without or with doxorubicin (DOX, 0.5 μM) for 4 hr. Cells were harvested and analyzed by immunoblotting for phospho-E6AP Ser-218 and α-tubulin. (B) HEK293 cells were treated without or with DOX (0.5 μM), or ATM inhibitor (KU55933, 10 μM), for 4 hr, and analyzed by immunoblotting. (C) HeLa cells were treated with hydroxyurea (HU, 10 mM) combined with ATM/ATR inhibitor (caffeine, 4 mM) or ATM inhibitor (KU55933, 10 μM), for 12 hours, as indicated. Cells were harvested and analyzed by immunoblotting.

Figure 1 Supplemental 1

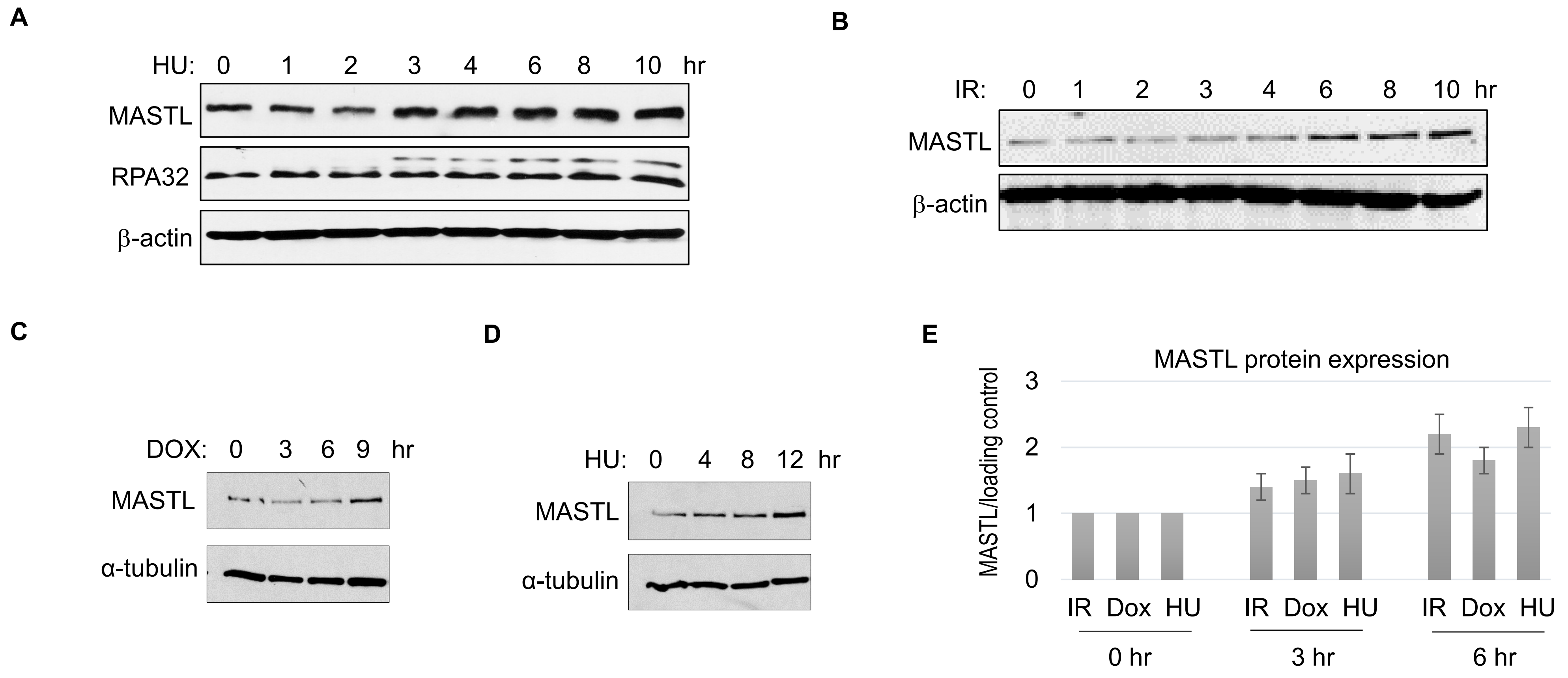

Figure 1 Supplemental 2

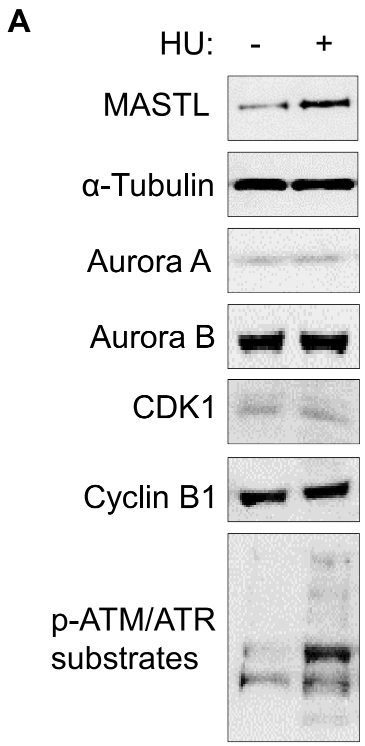


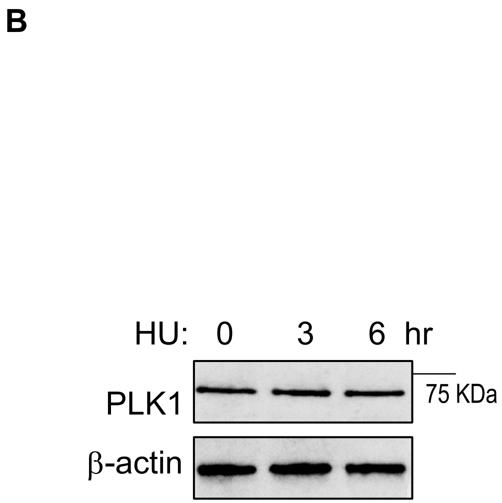

Figure 2 Supplemental 1

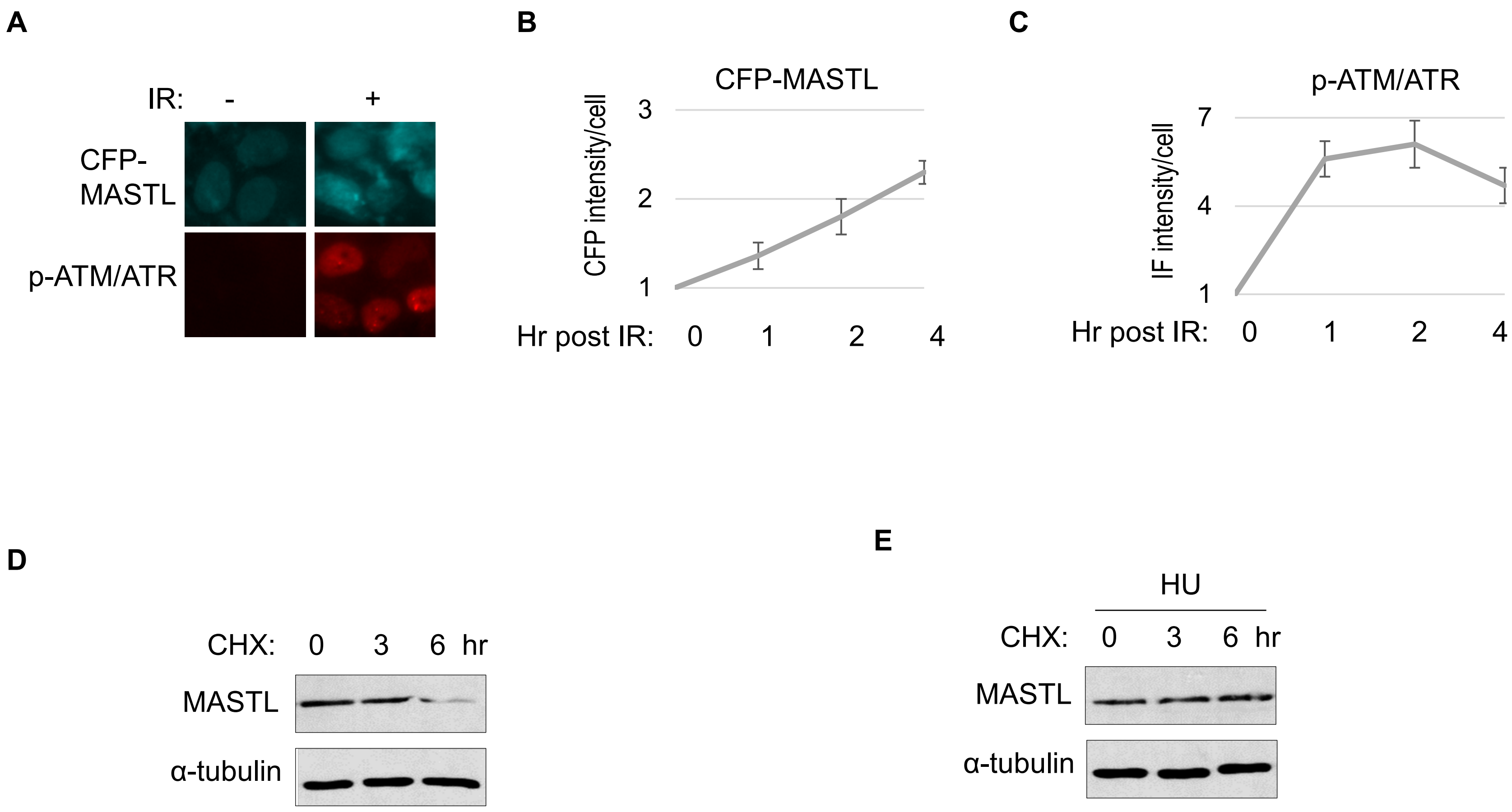

Figure 3 Supplemental 1

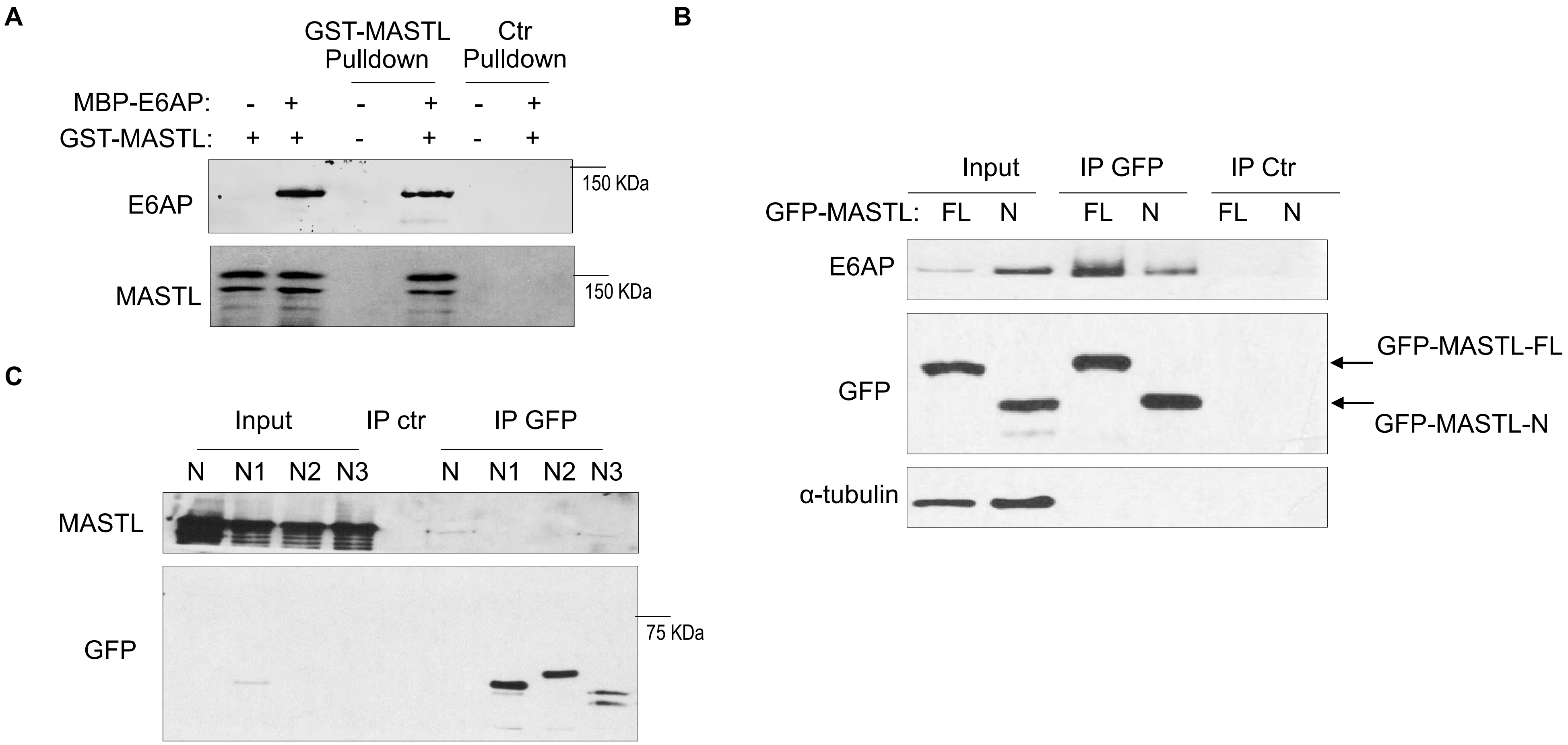

Figure 4 Supplemental 1

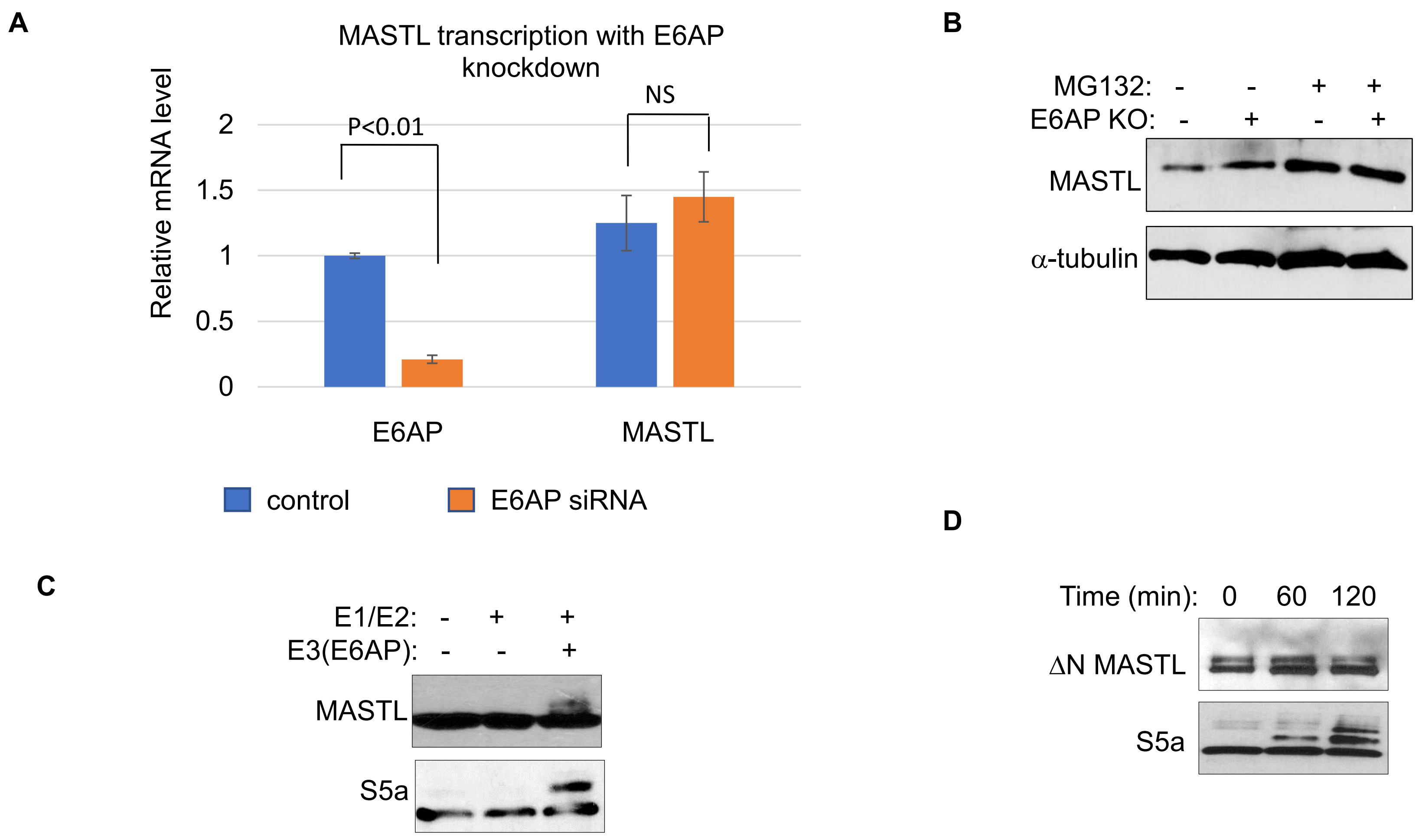

Figure 5 Supplemental 1

A

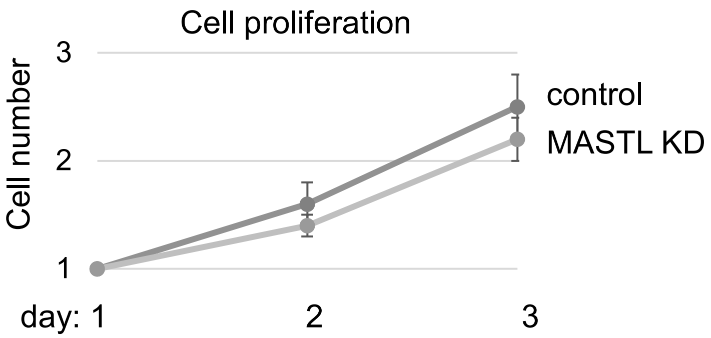


B

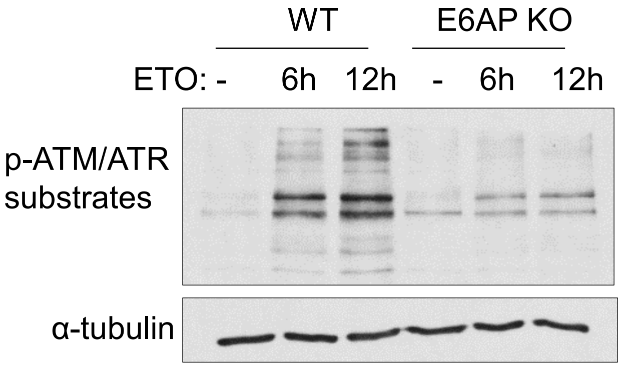

Figure 7 Supplemental 1

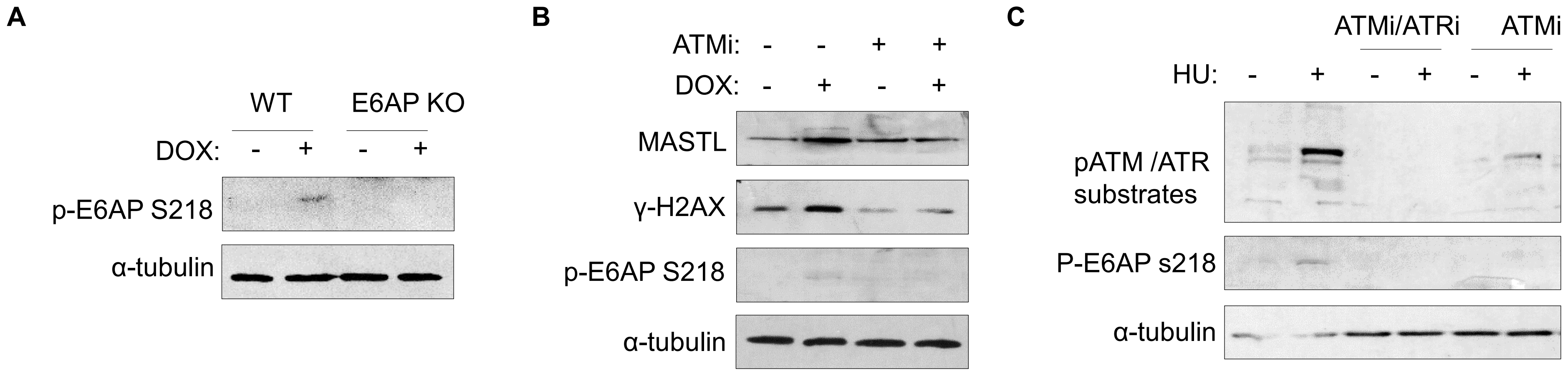
